# Evidence of population specific selection inferred from 289 genome sequences of Nilo-Saharan and Niger-Congo linguistic groups in Africa

**DOI:** 10.1101/186700

**Authors:** Julius Mulindwa, Harry Noyes, Hamidou Ilboudo, Oscar Nyangiri, Mathurin Koffi, Dieudonne Mumba, Gustave Simo, John Enyaru, John Chisi, Martin Simuunza, Pius Alibu, Vincent Jamoneau, Annette Macleod, Bruno Bucheton, Christiane Hertz-Fowler, Issa Sidibe, Enock Matovu, for the TrypanoGEN Research Group, as members of The H3Africa Consortium.

## Abstract

**Background:** There are over 2000 genetically diverse ethno-linguistic groups in Africa that could help decipher human evolutionary history and the genetic basis of phenotypic variation. We have sequenced 300 genomes from Niger-Congo populations from six sub-Saharan African countries (Uganda, Democratic Republic of Congo, Cameroon, Zambia, Ivory Coast, Guinea) and a Nilo-Saharan population from Uganda. Of these, we analysed 289 samples for population structure, genetic admixture, population history and signatures of selection. These samples were collected as part of the TrypanoGEN consortium project [1].

**Results:** The population genetic structure of the 289 individuals revealed four clusters, which correlated with ethno-linguistic group and geographical latitude. These were the West African Niger-Congo A, Central African Niger-Congo B, East African Niger-Congo B and the Nilo-Saharan. We observed a spatial distribution of positive natural selection signatures in genes previously associated with AIDS, Tuberculosis, Malaria and Human African Trypanosomiasis among the TrypanoGEN samples. Having observed a marked difference between the Nilo-Saharan Lugbara and Niger-Congo populations, we identified four genes (*APOBEC3G*, *TOP2B*, *CAPN9*, *LANCL2*), which are highly differentiated between the two ethnic groups and under positive selection in the Lugbara population (_iHS -log p > 3.0, Rsb -log p > 3.0, Fst > 0.1 bonferroni p > 1.8x10e4).

**Conclusion:** The signatures that differentiate ethnically distinct populations could provide information on the specific ecological adaptations with respect to disease history and susceptibility/resistance. For instance in this study we identified *APOBEC3G* which is believed to be involved in the susceptibility of the Nilo-Saharan Lugbara population to Hepatitis B virus infection.

## Background

The African continent’s ethno-linguistic groups have been classified into four major families, Afro-Asiatic, Nilo-Saharan, Niger-Congo, and Khoisan [2]. The Afro-Asiatic which includes the Semitic, Cushitic, and ancient Egyptian languages, is spoken predominantly by northern and eastern African pastoralists and agro-pastoralists; the Nilo-Saharan, which includes the Central Sudanic and Eastern Sudanic (Nilotic) languages, is spoken predominantly by eastern and central Saharan pastoralists; the Niger-Congo languages are subdivided into the Niger-Congo A in West Africa and the Niger-Congo B or Bantu in Central, Eastern and Southern Africa [3, 4]. Fourteen ancestral population clusters, which correlate with shared cultural and linguistic affiliations, have been identified amongst these groups [5]. These 14 ancestral populations further subdivide into over 2000 ethnically diverse linguistic groups [6, 7].

The genetic diversity of ethno-linguistic groups can be used to study human evolutionary history and the genetic basis of phenotypic variation [5] and complement studies of African genotype variations [5, 8–10] that have contributed to the understanding of human origins and disease susceptibility markers. However, samples from sufficient individuals for population analysis have been sequenced from relatively few African populations. The 1000 genome project generated data from five Niger-Congo populations, The African Variome project added Afro-Asiatic populations and there have been small scale studies of the Khoisan hunter-gatherers [5, 8, 11, 12]. To date, no sequences of Nilotic populations have been published, although one previous study used 200,000 SNP loci to examine genetic diversity of the Nilo-Saharan speaking population of southern Sudan Darfurian and Nuba peoples [13]. In the present study we present the first genome sequences of a Nilo-Saharan population and genome sequences from six new Niger-Congo populations.

## Results

### Samples and sequencing

The samples used for this study were collected by the TrypanoGEN consortium and consisted of 300 individuals from 17 linguistic groups who were residents of Guinea, Ivory Coast, Cameroon, Democratic Republic of Congo, Uganda and Zambia (Table 1, Additional file 1: Table_S0). DNA was extracted from blood and genomes were sequenced on the Illumina 2500 platform at 10X coverage, except for the Zambia and Cameroon samples that were sequenced at 30X coverage.

**Table 1.** Table showing the Ethnic groups and number of individual from each Country that were used for Whole genome sequencing. The Ethnicity codes were obtained from the Ethnologue languages of the World catalogue [80].

| Country | Code | Ethnicity | Ethnologue code | No. of Samples |
| --- | --- | --- | --- | --- |
| Uganda | UNL | Lugbara | lgg | 50 |
|  | UBB | Basoga | xog | 33 |
| Zambia | ZAM | Soli/Chikunda | sby; kdn | 28 |
|  |  | Tumbuka | tum | 14 |
|  |  | Bemba | bem | 8 |
| Congo | DRC | Kimbala | mdp | 20 |
|  |  | Kingongo | noq | 30 |
| Cameroon | CAM | Bamilike | fmp | 6 |
|  |  | Mundani | mnf | 8 |
|  |  | Ngoumba | nmg | 12 |
| Ivory Coast | CIV | Baoule | bci | 11 |
|  |  | Gouro | goa | 21 |
|  |  | More | Moa | 12 |
|  |  | Senoufo | sef | 4 |
|  |  | Malinke | loi | 1 |
|  |  | Koyaka | kga | 1 |
| Guinea | GAS | Soussou | sus | 50 |

Following mapping and SNP calling, we identified approximately 34.1 million single nucleotide polymorphisms (SNPs) and 5.3 million insertion/deletion (Indel) polymorphisms (Table 2). We identified 2.02 million variants that did not have rsIDs and were considered ‘novel’. The SNPs had a transition-transversion ratio of 2.0 (Additional file 2: Figure S1), implying good quality SNP calls [14, 15]. Prior to population analysis, variants (SNPs and Indels) were filtered by removing loci with >10% missing data, Minor Allele Frequency (MAF) < 0.05 or Hardy Weinberg Equilibrium (HWE) P-value < 0.01. Eleven individuals with > 10% loci missing were removed from the original 300 individuals dataset (Table 2), leaving 289 samples for downstream analysis. An additional 504 samples from five African populations in the 1000 genomes project (Esan and Yoruba from Nigeria, Mende from Sierra Leone, Mandinka from Gambia and Luhya from Kenya), were included in some of our analyses.

**Table 2.** The number of SNPs and Indels obtained from the mapping and variant; alling pipeline. The SNPs were filtered for HWE, MAF and missing genotypes

| Pop | Samples<br>(WGS) | Samples<br>after<br>filtering | SNPs before<br>filtering | SNPs<br>after<br>filtering | Indels<br>before<br>filtering | Indels<br>after<br>filtering |
| --- | --- | --- | --- | --- | --- | --- |
| **CIV** | 50 | 50 | 18,780,913 | 16,066,827 | 3,069,408 | 1,583,594 |
| **DRC** | 50 | 49 | 19,188,537 | 16,449,696 | 3,146,802 | 1,626,826 |
| **GAS** | 50 | 47 | 18,831,834 | 16,075,002 | 3,063,080 | 1,579,352 |
| **UBB** | 33 | 32 | 17,671,306 | 14,987,699 | 2,889,915 | 1,426,646 |
| **UNL** | 50 | 44 | 18,986,243 | 15,598,629 | 3,130,979 | 1,536,490 |
| **CAM\*** | 26 | 26 | 17,183,994 | 14,579,603 | 3,283,543 | 1,539,459 |
| **ZAM\*** | 41 | 41 | 18,232,386 | 15,548,110 | 3,448,501 | 1,651,467 |
| **Total** | **300** | **289** | **34,116,333** | **30,591,165** | **5,336,622** | **3,166,196** |

\* Samples Sequenced at 30X coverage, rest of the samples were at 10X coverage. Identified 2,023,049 SNPS without rsIDs {Total SNPs (30,591,165) – SNPs with rsIDs (28,568,116)}

### Population stratification by Multiple Dimensional Scaling

Multiple Dimensional Scaling (MDS) implemented in Plink 1.9 software was used to help visualise genetic distances between samples (Figure 1). All TrypanoGEN samples clustered by country except those in Uganda, where the Nilo-Saharan Lugbara samples formed a distinct cluster from the Niger-Congo B Basoga samples. When the samples from the six TrypanoGEN and the four African 1000 genomes project countries were merged, five groups representing five major geographic groups were observed (Figure 1B): the Uganda Nilo-Saharan; East African Bantu speakers from Uganda and Kenya; Central African Bantu speakers from Cameroon, DRC and Zambia; Nigerian Niger-Congo A speakers (Esan and Yoruba); West African Niger-Congo A speakers from the Ivory Coast, Gambia, Sierra Leone and Guinea. The African and European samples were very distinct (Figure 1C). Since all samples except Ugandan Bantu and Nilo-Saharan clustered by country by MDS, samples were grouped by country for subsequent analyses except for the Uganda samples, which were grouped by both country and linguistic group.

**Figure 1.**
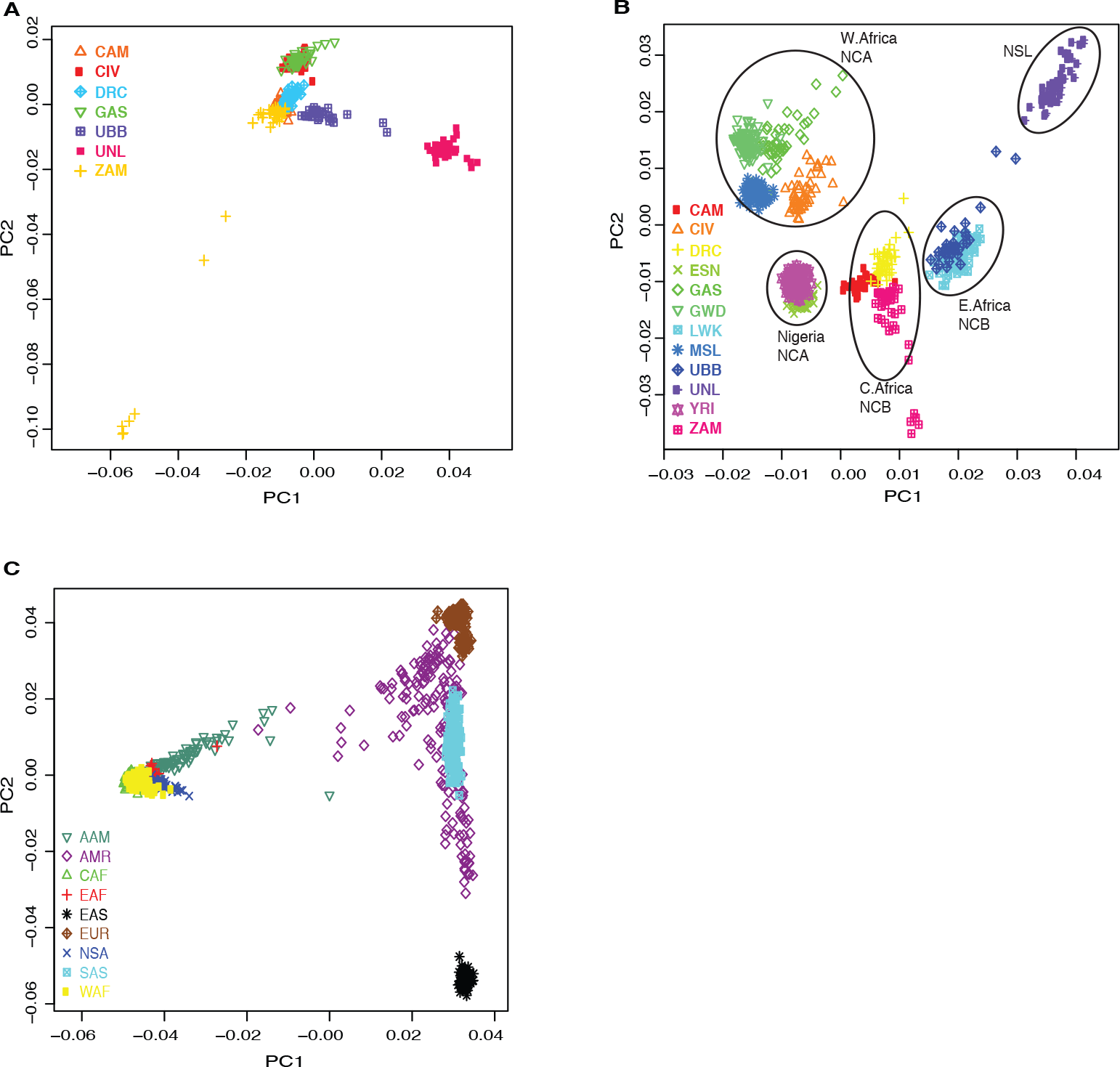
Multi dimensional scaling (MDS) analysis of the sequenced TrypanoGEN samples, Guinea (GAS), Ivory Coast (CIV), Cameroon (CAM), Democratic Republic of Congo (DRC), Uganda (Nilotics, UNL, Bantu, UGB) and Zambia (ZAM), (**A**); **B**, TrypanoGEN and selected 1000 genomes African samples Nigeria (ESN, YRI), Sierra Leone (MSL), Gambia (GWD), Kenya (LWK); The black circles indicate the clustering of the samples into the West African Niger-Congo-A, Nigerian Niger-Congo-A, Central African Niger-Congo-B, East African Niger-Congo-B and the Nilo-Saharan Lugbara (NSL); **C**, 1000 genomes samples from Africa and the rest of the world. AAM, African Americans; AMR, indigenous Americans; CAF, Central Africa; EAF, East Africa; EAS, East Asia; EUR, Europe; NSA, Nilo-Saharan; SAS, South Asia; WAF, West Africa;

### Population Admixture and differentiation

The amount of shared genetic ancestry within the samples was estimated using the Admixture software [16]. Admixture was run on 2-8 population clusters (K) in triplicate; with K=4, K=5 and K=6 having the lowest cross validation errors and hence the most probable numbers of ancestral components represented in the data (Additional file 2: Figure S2). At K=6 the Niger-Congo populations exhibited 17-60% admixture with minor ancestries, whilst the Ugandan Nilo-Saharan population had a platry 7% admixture with Niger-Congo ancestries (Figure 2A). At K4 one European and three ancestral African populations were observed, which corresponded to Nilo-Saharan, Niger-Congo-B (East African) and Niger-Congo-A (West African). At K5 a homogeneous group of seven samples emerged within the Zambia population with no admixture with other populations in our data set and were also outliers on the MDS plot (Figure 1B). These seven were recorded as Soli/Chikunda speakers, which are Bantu languages but they had no admixture at (K=5 and K=6) with the other speakers of this language group from Zambia or any other group included in this study, suggesting that they had a quite distinct ancestry. At K6, a major group appeared that contributed ancestral components to both East African Niger-Congo B and West African Niger-Congo A but did not correspond to any existing linguistic group.

**Figure 2.**
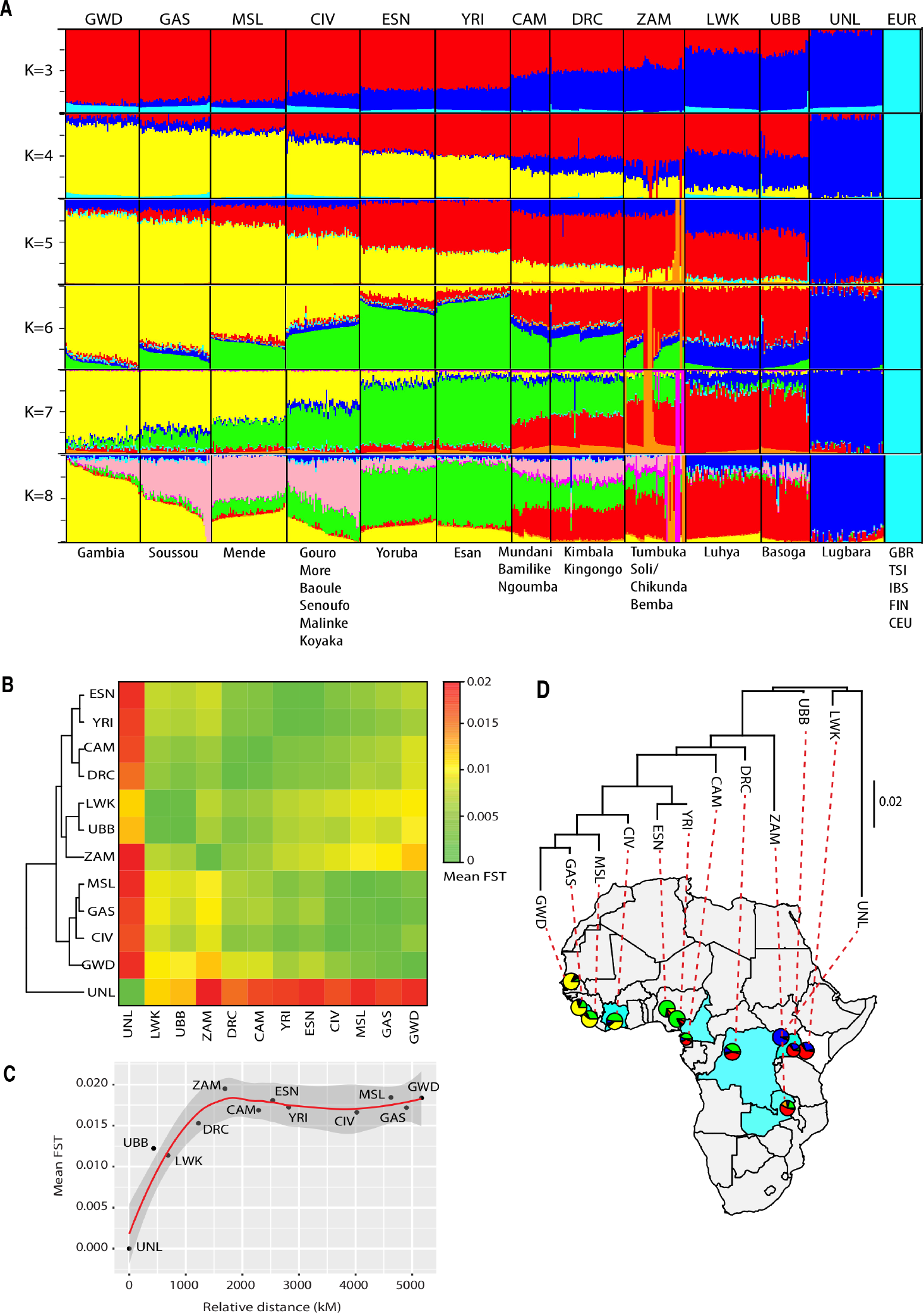
Genetic admixture and diversity between TrypanoGEN and selected 1000 genome populations. **A**. Admixture plot of the K populations of the TrypanoGEN, 1000 genome African and European populations. **B**. Heatmap of mean Fst between TrypanoGEN and 1000 genome African populations. **C**. Polynomial regression plot of the mean Fst against the relative geographical distance of the African Niger-Congo populations from the Uganda Nilotic population. **D**. Phylogeographic plot of the mean Fst distances on the Trypanogen populations and selected 1000 genome African populations; the pie charts represent the population sample size and admixture.

Genetic variation within the populations that are part of the TrypanoGEN project was estimated using the pairwise F_ST_ [17] (Figure 2B, Additional file 2: Figure S3). F_ST_ was relatively high between the Nilo-Saharan Lugbara samples and the African Bantu populations (Figure 2B) except the East African Basoga (population mean F_ST_ = 0.012) and Luhya (population mean F_ST_ = 0.011), presumably due to the 30% admixture of Nilo-Saharan origin within these populations. The pattern of the observed genetic variation was consistent with the relative geographic distance from the Nilo-Saharan population (Figure 2C). In addition, a classification based on the genetic distances between populations (F_ST_) showed clustering of populations by geographic region on the African continent (Figure 2D).

### Population size over time and timing of population isolation

The Multiple Sequentially Markovian Coalescent (MSMC) method was used to estimate population sizes over time and times at which populations became isolated (Figure 3). Effective population sizes (N_e_) were relatively stable at around 13,000 in all populations tested from 100 thousand years ago (kya) until about 50kya when they started to decline reaching a nadir of about 8,000 about 13kya coinciding with the dry period at the end of the last ice age (Figure 3A, Additional file 3: Table S1). All population sizes increased rapidly thereafter but the Niger-Congo populations increased to an N_e_ of around 200,000, whilst the Nilotic population only increased to 60,000. The Ugandan Bantu population was intermediate in N_e_ presumably due to admixture with the Nilotics. This post glacial population increase was briefly reversed in the Central and West African populations which suffered declines of 6-23% between 1500 and 750 years ago before recovering to even higher levels at the present time. This absolute decline in N_e_was not observed in the Ugandan Bantu population, although the growth rate declined. In the Nilotic population, a decline was observed at a later time point after 750 years ago.

**Figure 3.**
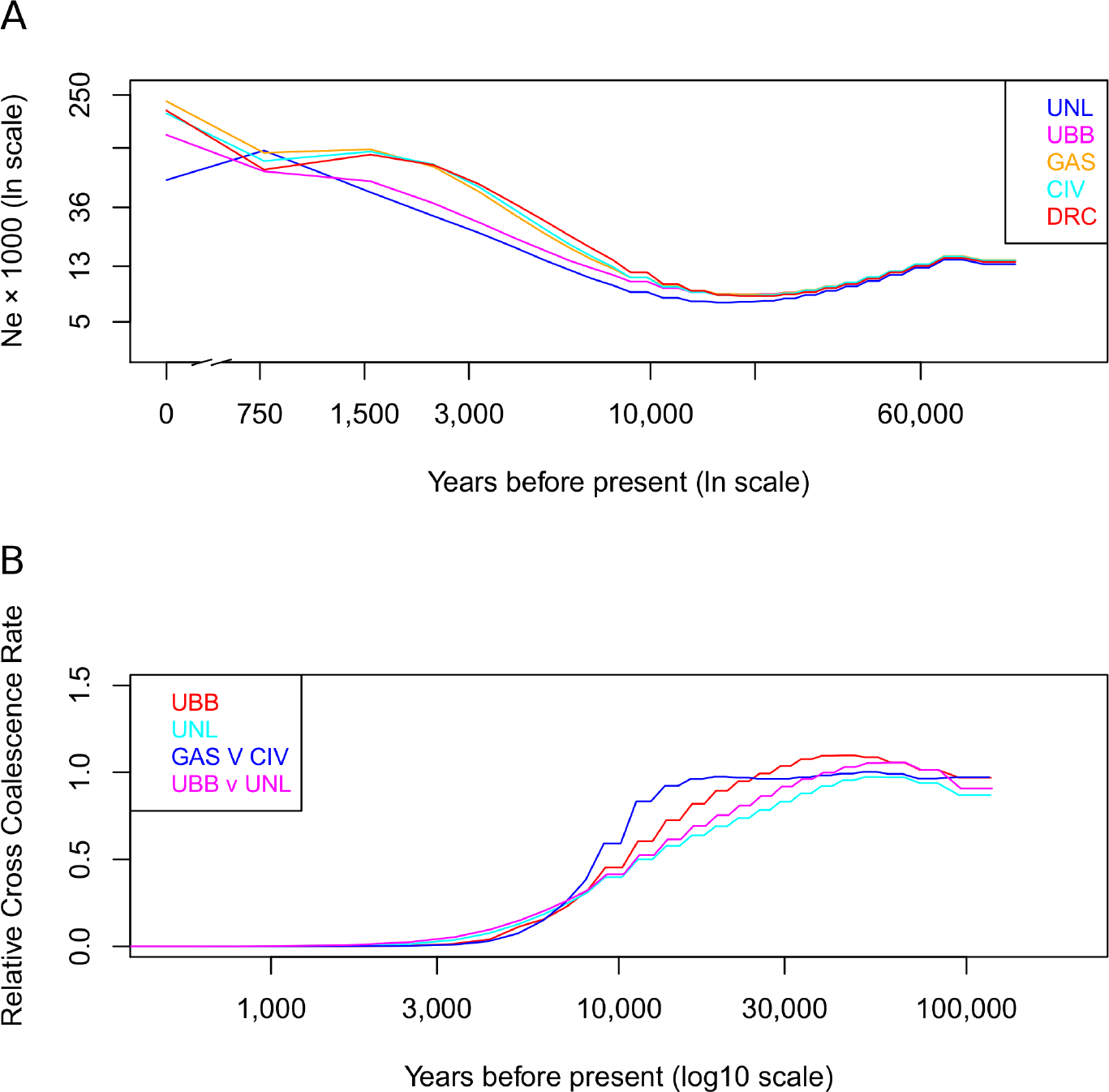
Population sizes and cross-coalescence rates compiled by MSMC. **A** Effective population sizes for each population since 75kya. The Ugandan Bantu and Nilotic populations have grown continuously but at a slower rate than the West and Central African populations since 13kya. These latter populations experienced declines of 6-23% between 1500 and 800 years ago. **B** Cross-coalescence rates for pairs of populations. At 1.0 populations are panmictic and at 0.0 there is no gene flow. The Guinea and Ivory Coast populations were panmictic until about 10 kya and then became separated by 3kya. Other populations appear to have separated more gradually but these may be confounded by admixture.

Population separation data is less clear and may be more sensitive to admixture (Figure 3B). The Guinea and Ivory Coast populations were the least admixed and appeared panmictic until about 10kya, and had become isolated by about 3kya. The Ugandan Bantu and Ugandan Nilotic appeared to begin separating from other populations about 23 and 47kya, respectively and became isolated about 3kya but these estimates may be confounded by admixture.

### Genome-wide screen for extended haplotypes under selection

#### Signatures within population

In order to identify alleles under selection pressure, we used the within population Extended Haplotype Homozygosity (EHH) test [2, 18]. Similar patterns of loci with extreme positive and negative iHS scores were observed across all groups (Additional file 4: Figure S4A). The iHS values for all groups had an approximate normal distribution (Additional file 4: Figure S4C) implying that the sizes of iHS signals from different SNPs in all the populations were comparable [3, 4, 19]. The mean number of loci with extreme positive and negative iHS score (-log p > 3) from all groups was 8,984, Guinea had the largest number of loci with extreme iHS score (11,401) and Zambia had the least (5,570) (Table 3, Additional file 5: Table S2). These extreme loci were classified by the Ensembl annotation of the nearest gene. Approximately 34% of these annotations were for protein coding genes; a mean of 3,058 SNPs in protein coding genes per population were associated with extreme iHS scores. Some protein coding genes with extreme iHS SNP loci were shared between different countries whereas some occurred only in a single country population (Additional file 5: Table S2, sheet ‘ALLpop.protein_coding’). We observed strong iHS signatures in genes that have been previously identified in other African populations as being under strong selection [5, 8, 19, 20]. These included ***SYT1***, a synaptosomal protein implicated in Alzheimer’s disease [6, 7, 21] that was found in all country populations; ***LARGE*** a glycosylase involved in Lassa fever virus binding [5, 22] (Zambia, Cameroon, Ivory Coast); ***CDK5RAP2*** a microcephaly gene controlling brain size [5, 8–10, 23] (Ugandan Bantu); ***NCOA1***, a transcriptional coactivator associated with Lymphoma (Guinea, Ivory Coast, DRC); ***SIGLEC12***; involved in immune responses [5, 8, 11, 12, 24] (Zambia, Cameroon). Using the DAVID annotation database [13, 25] we observed that all of the country populations had strong signals that have been implicated in communicable diseases such as HIV/AIDS, Malaria and Tuberculosis which have high prevalence in Africa [14, 15, 26] (Table 4), suggesting an adaptive role of these genes to infection.

**Table 3.** Extreme iHS loci that overlap with the UNL population

| Pop | Extreme iHS<br>SNPs ($-\log p > 3.0$) | Extreme iHS SNPS<br>associated with protein<br>coding genes | Extreme iHS SNPs<br>overlapping with<br>UNL |
| --- | --- | --- | --- |
| **UNL** | 8454 | 2613 | 2613 |
| **UBB** | 9617 | 3326 | 512 |
| **DRC** | 10037 | 3790 | 535 |
| **ZAM** | 5570 | 1990 | 86 |
| **CIV** | 10129 | 3541 | 534 |
| **CAM** | 7686 | 2597 | 82 |
| **GAS** | 11401 | 3741 | 382 |

**Table 4.** DAVID [25] analysis of Genes that are highly selected within TrypanoGEN population and associated with HIV, Tuberculosis, and Malaria. The Fisher’s exact test *P*-values indicate significant gene enrichment in the associated disease

| Gene | Chr | Populations affected | Associated Disease | p value | Reference |
| --- | --- | --- | --- | --- | --- |
| *HLA-DRB1* | 6p21.32 | ZAM, CAM, CIV, DRC, UBB | HIV/TB/Malaria | 1.63E-09 | [81, 82] |
| *NLRP1* | 17p13.2 | ZAM, CIV, DRC, GAS, UBB | HIV | 2.42E-07 | [83] |
| *VPRBP* | 3p10.6 | UBB, CIV, DRC, GAS | HIV | 2.42E-07 | [84] |
| *TRIM5* | 11p15.4 | UNL, CAM, GAS | HIV | 7.30E-07 | [85] |
| *ANKRD30A* | 10p11.21 | DRC, CIV | HIV | 2.42E-07 | [86] |
| *HLA-A* | 6p22.1 | ZAM, CAM | HIV/TB | 4.70E-06 | [87] |
| *HLA-DQA1* | 6p21.32 | UBB, DRC | HIV/TB | 4.70E-06 | [87] |
| *HLA-DQB1* | 6p21.32 | UBB, DRC | HIV/TB | 4.70E-06 | [87] |
| *KIR3DL1* | 19q13.42 | UNL, CIV | Malaria | 1.63E-09 | [88, 89] |
| *CD36* | 7q21.11 | UNL, CAM, CIV | Malaria | 1.55E-06 | [90] |
| *DDC* | 7p12.2 | UBB, DRC, GAS | Malaria | 1.63E-09 | [91] |
| *HBE1* | 11p15.4 | UBB, CAM, DRC | Malaria | 5.48E-07 | [92] |
| *ADCY9* | 16p13.3 | UNL, CIV | Malaria | 5.48E-07 | [93] |

Having collected samples from Human African Trypanosomiasis (HAT) endemic regions, we identified signatures that have been implicated in trypanosome infection. These signatures were observed in genes overlapping the KEGG calcium signalling pathway [16, 27, 28] that were previously identified mainly from mouse studies [29]; ***F2RL1*** (Guinea, Ivory Coast), ***GNA14*** (Zambia), ***GNAQ*** (Cameroon), ***GNAL*** (Guinea, Cameroon), ***GNAS*** (Zambia). The calcium-signalling pathway regulates permeability of the blood brain barrier to trypanosome parasites during central nervous system (CNS) disease [30]. In addition, we observed signatures in genes overlapping the Mitogen-activated protein kinase MAPK pathway ***MAPK1*** (Cameroon), ***MAPK10*** (Ugandan Nilo-Saharan, DRC, Ugandan Bantu), ***MAPK9*** (Zambia); which is targeted by trypanosomatids in order to modulate the host’s immune response [17, 31]. These host signalling pathways have been shown to play a role in host immunity against trypanosome infection in mice and cattle [32–34].

#### Signatures unique to Nilo-Saharans

In order to determine which signatures are unique to the Nilo-Saharan Lugbara, we first ascertained which extreme iHS loci (-log p > 3) were common to the Nilo-Saharan and one or more Niger-Congo groups. We observed that approximately 15% of the protein coding gene associated extreme iHS SNPs of the Ugandan Bantu, DRC, Ivory Coast and Guinea groups were common with the Nilo-Saharan group, whereas Cameroon and Zambian groups had 2.7% in common (Table 3, Additional file 4: Figure S4B). We identified 149 extreme SNPs associated with protein coding genes and were unique to the Uganda Nilo-Saharan (Additional file 6: Table S3). Using the PANTHER Gene ontology database [35], we observed that these unique genes were mainly enriched for cellular and metabolic process proteins (approximately 50.8%) (Additional file 7: Figure S5). Amongst these were SNPs associated with genes that have also been shown by other studies to be under positive selection, including ***APOBEC3G***, which is involved in innate anti-viral immunity [36, 37], with demonstrated protective alleles against HIV-1 in Biaka and Mbuti pygmies of Central African Republic and DRC respectively [38]; ***IFIH1*** (also called ***MDA5***), a cytoplasmic RNA receptor that mediates antiviral responses by activating type I interferon signalling [39] but is also implicated in protection against type 1 diabetes ([40, 41]; ***OR2L13***, an olfactory receptor involved in activation of signal transduction pathway for odorant recognition and discrimination [42], and is associated with Diabetic nephropathy in African Americans [43].

#### Nilo-Saharan versus Niger-Congo cross population signatures

There were 299 SNPs with high F_ST_ (above 99^th^ percentile) and XPEHH (Rsb -log p > 3) in the regions of protein coding genes that were also highly differentiated between the Nilo-Saharan and Niger-Congo populations (Additional file 8: Table S4). We then compared SNP loci with derived alleles that are unique to the Nilo-Saharan Lugbara and occur in highly differentiated genes (extreme Rsb, high F_ST_) between the Nilo-Saharan and Niger-Congo groups. From this we identified 12 genes (Table 5, Additional file 9: Figure S6D) including the ***APOBEC3G*** gene, that are highly differentiated between the Nilo-Saharan Lugbara and Niger-Congo groups (mean F_ST_ 0.11, Rsb -log p = 4.1). ***APOBEC3G*** also contains the SNP rs112077004, which was observed to be under positive selection in the Nilo-Saharan Lugbara (Figure 4, Additional file 10: Figure S7, Additional file 11: Table S5).

**Table 5.**
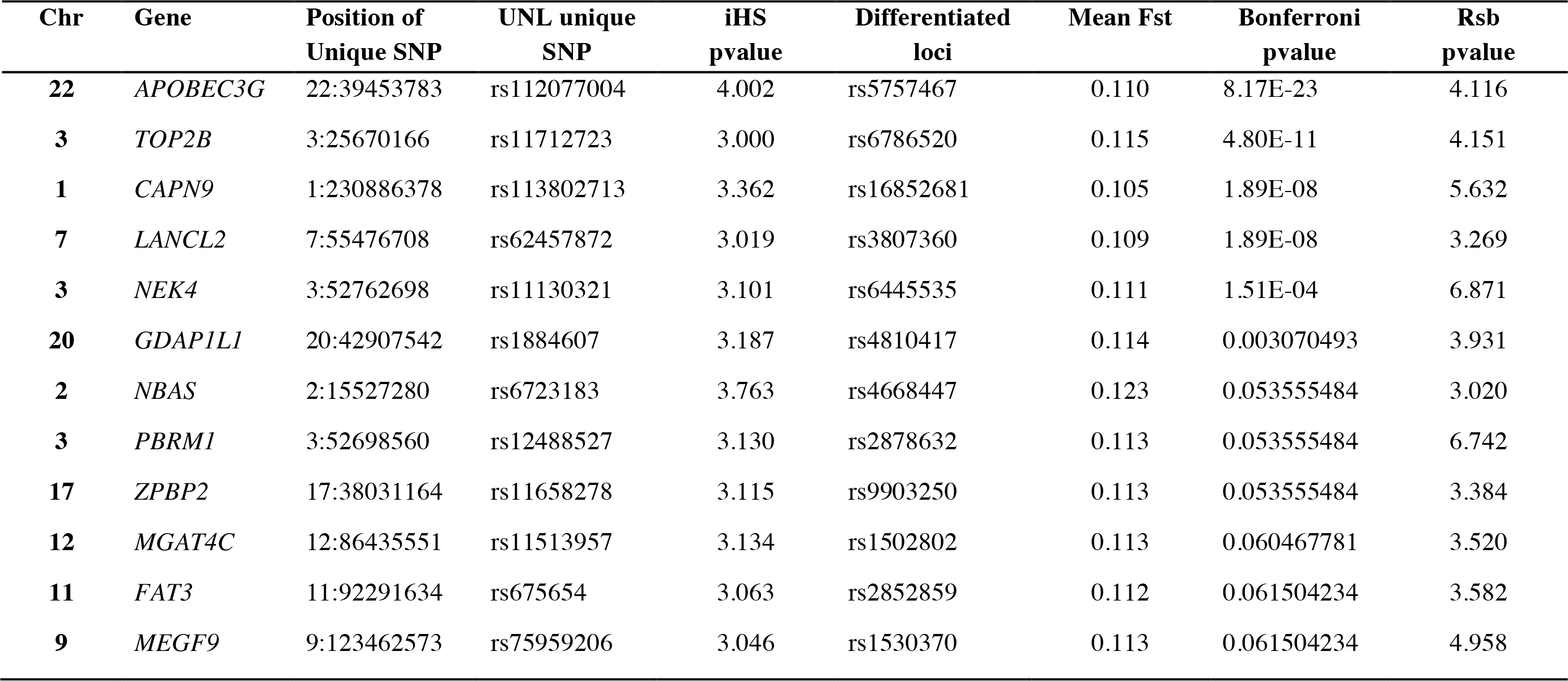
Genes that are highly differentiated between the Nilo-Saharan and Trypanogen Niger-Congo populations that contain SNPs unique to UNL population

**Figure 4.**
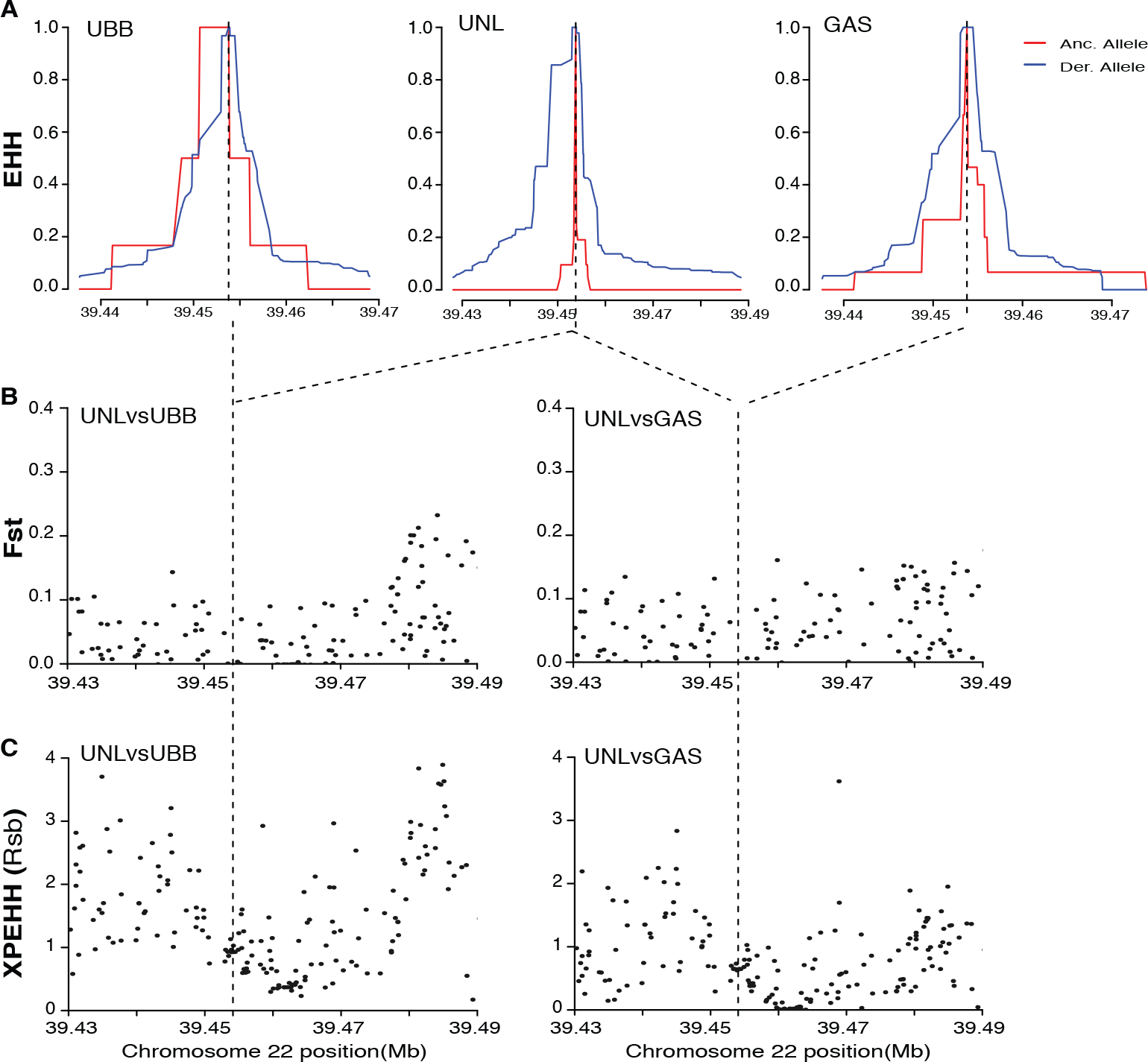
Illustration of signatures unique to the Uganda Nilotic population. Signal of positive selection within the *APOBEC3G* gene on Chromosome 22 at the rs112077004 loci of the Uganda Nilo-Saharan Lugbara population, in comparison with the Niger-Congo B populations of Uganda (UGB) and Niger-Congo A population of Guinea (GAS). **A**. The calculated site specific extended haplotype homozygosity (EHH) within a population. **B**. Between population Fst analysis. **C**. Across population (XPEHH) analysis.

## Discussion

We have analysed the genomes of 289 individuals from seven populations from six Sub-Saharan Africa Countries, investigating their admixture profile, demographic histories and signatures of selection that differentiate the major linguistic groups. The MDS analysis identified five major clusters: Nilo-Saharan, two Niger-Congo A groups from Nigeria and West Africa and two groups of Niger-Congo B (Bantu speakers) from Central and East Africa, which were consistent with previous studies [5, 8, 44]. The samples represented three of the five major linguistic groups in Africa, omitting the Afro-Asiatic and Khoisan speakers. Afro-Asiatic speakers are found across North and North-East Africa in regions adjacent to Nilo-Sharan and Bantu speakers. Afro-Asiatic reference populations were not included in this study and we therefore could not detect any admixture from this source. A SNP genotype based analysis of Nilotic populations indicated that Nilotic populations only contain a trace of Afro-Asiatic ancestry and therefore our observations on East African populations may not be significantly limited by the absence of Afro-Asiatic data [13].

Linguistic analyses suggest that Niger-Congo speaking hunter-gathers originated from the Kordofanian speakers of the Nuba mountains of Sudan and then traversed the Sahel to Mali (Figure 5). They colonised the coast from Senegal to Nigeria and Cameroon over several thousand years, forming multiple linguistic groups. The Bantu (Niger-Congo-B) speaking people emerged as another linguistic group amongst the greater than 60 Niger-Congo-A groups in the Nigeria/Cameroon region about 3,000 years ago. Bantu speaking peoples then spread South-East along savannah corridors through the Congo basin and emerged in the Great Lakes region and spread North to the Lake Victoria region and South down the East Side of Africa [10, 45, 46]. This rapid expansion is believed to have been enabled by the development of agriculture and later enhanced by the acquisition of iron tools [5].

**Figure 5.**
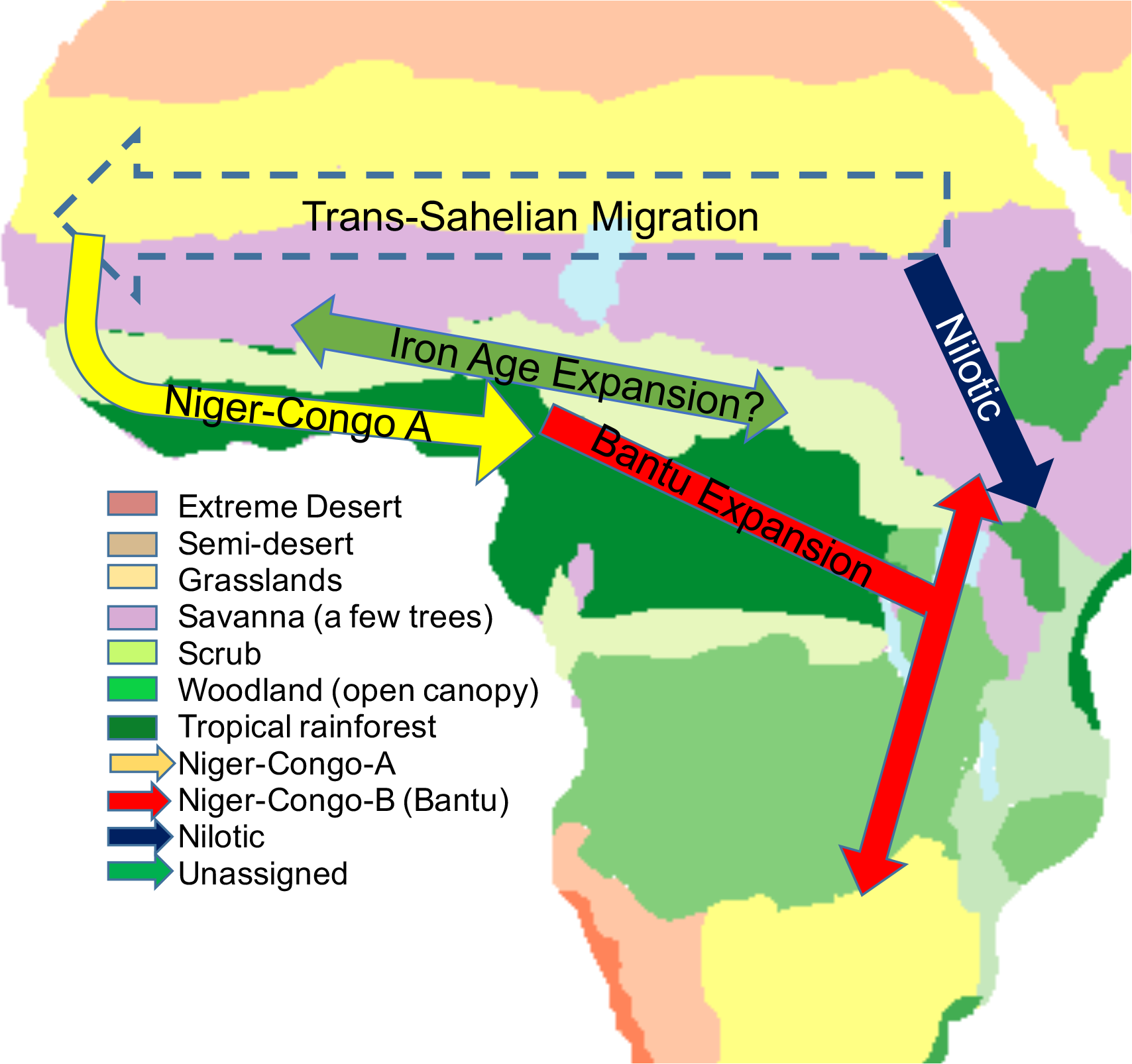
Migrations of Niger-Congo speakers. Map colours show vegetation coverage approximately 10kya [94]. Colours for linguistic groups as for Figure 2. Blue Nilo-Saharan; Yellow, Niger-Congo A; Red, Niger-Congo-B (Bantu); Green putative expansion of an ancestral group out of modern Nigeria. Blue dotted arrow, suspected route of proto-Niger-Congo-A speakers from Nuba mountains of Sudan to Senegal across the Sahel [95] when it was much wetter than at present.

Our admixture analysis at K=4 was consistent with this linguistic history and recent genetic analyses [8, 10] with three African ancestral allele clusters (AAC) which can be interpreted as representing Niger-Congo A languages in West Africa, Niger-Congo B (Bantu) in Central and East Africa and Nilo-Saharan in Northern Uganda. The Niger-Congo-A speakers in extreme West Africa appear to have approximately 10% Nilo-Saharan ancestry and this declines towards the East. On the other hand, the Bantu speakers are a mix of Niger-Congo-A and a distinct putative Bantu ancestral cluster that it at highest frequency in Nigeria and Cameroon, despite the Nigerian Yoruba and Esan not being Bantu languages. The Niger-Congo-A component is displaced by a Nilotic component with easterly latitude whilst the “Bantu” component remains constant. At K=5 a small AAC of 7 Bantu speakers from Zambia emerges, who evidently have a genetic heritage that does not match their self-declared linguistic affiliation, and are of unknown descent. At K=6 a fourth major African AAC appears (green in Figure 2) with strongest representation in the Nigerian Yoruba and Esan then tapering off east and west into Central and West Africa. This does not correspond to any linguistic group and displaces the Niger-Congo-A ancestry observed at K=4 and 5, to the east of Nigeria and Niger-Congo-B (Bantu) in Nigeria and to the West. This ancestral cluster could represent a secondary movement out of Nigeria of migrants who adopted their host’s language. One possible driver for such a migration, if it occurred, was the development of iron smelting which may have originated in Nigeria about 2,500 years ago [47]. Irrespective of the true number of ancestral allele clusters there is evidence of back migration of people with Bantu ancestral alleles into West Africa as has been observed before [44]. Interestingly this migration to the west was not accompanied by language expansion as it was to the east.

Regarding population history, the estimates of current N_e_obtained from our data with MCMS (Figure 3A) of around 200,000 in West and Central Africa and 57,000-125,000 in East Africa (Supplementary Table S6) was consistent with previous observations on other African samples using the same method [48][12], but our larger dataset provides higher resolution at recent time points than a previous analysis [12]. The faster growth in the Niger-Congo A and B than the Nilotic populations appears to predate the Bantu expansion. The Niger-Congo A population was believed to be expanding through West Africa as the climate became wetter after 10kya, consistent with the separation times between the Guinea and Ivory Coast populations observed on the Cross-Coalescence Plot (Figure 3B). The Nilo-Saharan population developed a pastoralist economy probably after 6kya but their expansion into the tsetse belt may have been delayed by trypanosomiasis and other diseases until the cattle developed tolerance [49–51]. The effective population size did not grow as fast as that of the Niger-Congo-A populations. The brief population decline dated at ~1340CE by MSMC coincides with the timing of the Black Death (1343-1353), however time resolution is low and the decrease was only observed at a single time point. There is evidence of abandonment of multiple large settlements throughout West Africa around the time of the Black Death and there is speculation that this was caused by the plague [52]. The decrease at this time appears to have impacted the West and Central African Niger-Congo but not the East African populations. Both Bantu and Nilotic populations in East Africa were cattle keepers and pastoralists to varying degrees [51] and the concomitant lower population density and mobile lifestyle may have made them less vulnerable to plague than the more settled and urbanised West Africans. The more recent decline in the Nilotic Lugbara effective population size is unexplained, but the catastrophic Rinderpest outbreak in the 1880’s and 1890’s that killed up 90% of indigenous cattle, which lead to the depopulation of the East African savannahs and may have ended the dominance of the Nilotic speaking Maasai over the Bantu Kikuyu could have been a contributory factor [53].

The Cross-Coalescence plots for comparison between populations other than the Guinea and Ivory Coast Niger-Congo-A show long periods of separation (not shown). This is not consistent with previous observations [12] or with the Ugandan Bantu populations having separated from Niger-Congo-A populations even more recently than the separation between Guinea and Ivory Coast populations, and is presumably due to the extensive admixture with the Nilotics observed in this population. The Central African cross-coalescence data also indicated older separation times than linguistic evidence suggests (not shown) and although there was less evidence of admixture in this population, these data should be treated with caution.

We identified evidence of selection in genes that have previously been associated with HIV/AIDS, Tuberculosis and Malaria. Given the high prevalence of lethal infections on the continent [26], the finding of positive natural selection at disease susceptibility loci is not surprising. However not all these genes occurred in all the populations, demonstrating spatially varying selection probably due to differing environmental pressures [54, 55]. We looked for signatures of selection in genes and pathways that are implicated in trypanosome infection, including the calcium signalling pathway [29, 30], the MAPK pathway [31–34], and ***HPR***, ***APOL1***, ***IL6***, ***HLAG***, genes [56–61] (Additional file 12: Figure S8, Additional file 13: Table S6). However we only found evidence for selection for the calcium signalling and MAPK pathway genes. This suggests that Trypanosomiasis may have had a selective force in these populations.

In order to determine signatures of selection unique to the Nilo-Saharan Lugbara population, we used a combination of linkage disequilibrium-based method (iHS and Rsb) and population differentiation based method (F_ST_) [62]. Using this approach we identified 12 loci associated with coding genes, which are unique to the Nilo-Saharan Lugbara and highly differentiated from the Niger-Congo population. Among these was the variant associated with ***APOBEC3G*** that demonstrated significant positive selection in the Nilo-Saharan Lugbara population. This protein is involved in viral innate immunity [63], by inducing a high rate of dC to dU mutations in the nascent reverse transcripts leading to the degradation of the viral genome [36, 37]. The Lugbara have relatively low prevalence of HIV (4%) in comparison to the Basoga (6.4%) and Baganda (10.7%) Bantu groups of Uganda but relatively high prevalence of Hepattis B suggesting that either ***APOBEC3G*** has different effects on each of these viruses [64–69].

We also identified the missense variant rs10930046 (T/C) located in the ***IFIH1*** CDS, which was unique to the Nilo-Saharan Lugbara and highly selected (iHS -log p 3.264). This gene is associated with up regulation of type I interferon signalling occurring in a spectrum of human diseases [39] and is believed to be involved in the suppression of Hepatitis B viral replication [70]. Being a non-synonymous variant, rs10930046 could alter the functioning of IFIH1 and thus increase susceptibility to HBV in the Lugbara population, something that could be tested by a candidate gene study for DNA virus infections. Northern Uganda is considered to have one of the highest prevalence of Hepatitis B virus in the world [71] which has perhaps resulted in a unique adaption of the Nilo-Saharan Lugbara population to infection.

## Conclusion

We have incorporated a Nilo-Saharan population into the analysis of genomic sequences of Niger-Congo populations for the first time and show extensive admixture between Nilo-Saharan ancestry and Niger-Congo B (Bantu) populations. We show evidence for signatures of selection within the Nilo-Saharan population in genes associated with infectious diseases that have different prevalence from surrounding Bantu (Niger-Congo B) populations.

## Methods

### Sample collection

The samples used for this study are part of the TrypanoGEN biobank [72], which describes ethics approval, recruitment, sample processing and the meta data collected. The ethical approval for the study was provided by the national ethics councils of the TrypanoGEN consortium countries involved in the sample collection.

The sampled populations were assigned a 3 letter code with the following criteria: Country name first letter, was used as the first letter of the code; For countries where more than one linguistic group was sampled, we used abbreviations of the country name, ZAM-Zambia, CAM-Cameroon, CIV-Ivory Coast, DRC-Democratic republic of Congo; For countries where a homogeneous linguistic group was sampled, we used the Country, language group and language, UNL-Uganda Nilotic Lugbara, UBB-Uganda, Niger-congo B (Bantu), Basoga, GAS-Guinea, Niger-Congo A, Soussou.

Peripheral blood was collected from the participants at the field sites, frozen, and transported to reference laboratories from where DNA extraction was carried out using the Whole blood MidiKit (Qiagen). The DNA was quantified using the Qubit (Qiagen) and approximately 1 μg was shipped from each country for sequencing at the University of Liverpool, UK except for Cameroon and Zambia whose DNA was shipped to Baylor College, USA.

### Sequencing and SNP calling

The whole genome sequencing libraries were prepared using the Illumina Truseq PCR-free kit and sequenced on the Illumina Hiseq2500. The samples from Guinea, Cote D’Ivoire, Uganda and DRC were sequenced to 10x coverage at the Center for Genomic Research at the University of Liverpool. The samples from Zambia and Cameroon were sequenced to 30X at the Baylor College of Medicine Human Genome Sequencing Center.

The sequenced reads were mapped onto the 1000 genomes project human_g1k_v37_decoy reference genome using BWA. The SNP calling on all the samples was done using the genome analysis tool kit GATK v3.4. The SNPs were then filtered by; a) removing loci with > 10% missing SNP, b) removing individuals with > 10% missing SNP loci and c) removing loci with Hardy Weinberg P value < 0. 01. In addition, loci with MAF < 0.05 were also removed for the population stratification and Admixture analysis. Variant annotation was done using snpEff (www.snpeff.sourceforge.net).

### MDS analysis

Population stratification was done using Multi dimensional scaling tool in Plink 1.9 and R v 3.2.1 tools. For this analysis, in addition to the filtering mentioned above, SNP loci less than 2000bp apart were removed in order to reduce the linkage disequilibrium (LD) between adjacent SNP. MDS analysis was carried out for (i) all TrypanoGEN data, (ii) all TrypanoGEN data plus African 1000 genome data, (iii) all TrypanoGEN data including 50 European and all African 1000 genome data excluding African Caribbean in Barbados (ACB) and African Southwest USA (ASW) populations.

### Population Admixture

The population ancestry of each individual was obtained using Admixture 1.23 [16] on the filtered PLINK .bed files on the same TrypanoGEN, one thousand genome African and European population data sets analysed by MDS. Admixture was run on K1 to K8 with three replicates for each run. The Admixture plots were drawn using the R tool ‘strplot’ [73].

### Genetic diversity: F_ST_

The genetic diversity due to difference in allele frequency among populations was analysed by the inter-population Wright’s F_ST_ [17] in PLINKv1.9. The F_ST_ estimates were made between TrypanoGEN (UNL, UBB, DRC, CIV, GAS) and one thousand genome African (LWK, YRI, ESN, MSL, GWD) populations. The F_ST_ dendrogram was generated using Fitch in Phylip3.685 [74]. The geographic distance matrix between populations was calculated based on their global position system (GPS) coordinates [75].

### Population History

Population sizes and divergence times were calculated using MSMC [48] on data phased with Beagle [76]. Since PCA and Admixture analysis had indicated little difference between linguistic groups in each country with the exception of the Ugandan Bantu and Nilotic populations, samples from each country with highest coverage were analysed together except for Uganda where Bantu and Nilotic samples were analysed as separate populations. For population size estimates, output from 3 independent runs each using 8 different haplotypes were combined. Using 8 haplotypes rather than 4 gives higher resolution at more recent time points. For estimates of relative cross coalescence rate, three replicate runs were done, each using 2 different samples (4 haplotypes) from each pairwise comparison between populations. Results presented are the means of the replicates.

### Signatures of selection

The estimation of haplotypes was carried out by first Phasing of the genotyped SNPs using SHAPEIT v2.2 software [77]. The extended haplotype homozygosity (EHH) was then analysed using the R software package *rehh* [78]. Two main EHH derived statistics were calculated from the phased haplotype data, that is, intra-population integrated haplotype Score (iHS) [19] and inter-population Rsb [79]. Bedtools v2.26.0 was used to identify the intersection of the iHS, Fst and Rsb loci.

## Declarations

### Ethical approval

The ethical approval for the study was provided by the national ethics councils of the TrypanoGEN consortium countries involved in the sample collection which are: Uganda (HS 1344), Zambia (011-09-13), Democratic Republic of Congo (No 1/2013), Cameroon (2013/364/L/CNERSH/SP), Cote d’Ivoire (2014/No 38/MSLS/CNER-dkn), and Guinea (1-22/04/2013). All the participants in the study were guided through the consent forms, and written consent was obtained to collect biological specimens.

### Consent of publication

Informed consent was provided by the study participants, for sharing and publishing their anonymised data.

### Availability of data and material

The datasets generated and/or analysed during the current study are available from the corresponding author on reasonable request. The sequenced data will be submitted to the EGA by H3ABionet under the study accession number EGAS00001002602.

### Fundind

The study was under the TrypanoGEN project, funded by the Wellcome Trust (099310/Z/12/Z). The funders had no roles in the design of the study and collection, analysis, and interpretation of data and in writing the manuscript.

### Author’s contributions

EM, AM, IS, BB, CHZ conceived the study; HI, MK, DM, GS, JE, JC, MS, PA, collected samples; JM and HN analysed the data, interpreted output and wrote the manuscript; All authors read and approved the final manuscript.

## Acknowledgements

The authors would like to acknowledge the study participants who donated their specimens, the personnel involved in the community engagement and coordinating sample collection and processing, the National sleeping sickness control programmes of the participating Countries. Z Lombard (University of Witwatersrand) and D Adeyemo (NHGRI) for facilitating sequencing of samples from Zambia and Cameroon at Baylor College of Medicine. The H3ABionet for training and support on data analysis. Also Fiona Marshall and Rebecca Grollemund for helpful discussions of African History.

## Additional files

**Additional file 1:** Table S0.xlsx. List of participant samples sequenced indicating their sequencing ID, country of origin, language and sex of the individual.

**Additional file 2:** Figure S1-S3.pdf: Figure S1. Variant annotation of the population whole genome sequence (WGS) data using snpEff. **a**. count of number of substitutions for each base combination, **b**. count of number of Insertion-Deletions with corresponding InDel length of affected loci **c**. Count of number of sites with Transition-Tranversion ratios with the corresponding allele frequency, **d**. Percentage count of the variant annotation classifications. Figure S2. Plot of the coefficient of variation (CV) error verses the K population (number of clusters). The K population with the lowest CV error is considered as the most appropriate K for the admixture. Figure S3. Bar plot of the mean F_ST_ between TrypanoGEN populations.

**Additional file 3:** Table S1.xlsx. Table showing MSMC output of effective populations sizes at different time points

**Additional file 4:** Figure S4.pdf. Extended haplotype homozygosity analysis of the TrypanoGEN populations for signatures of natural selection. **A**. Manhattan plot of the Intra-population integrated haplotype score (iHS) showing SNPs with extreme positive (> 2.5) and extreme negative (< −2.5). **B**. Bar plot showing the number of SNPs associated with protein coding with iHS > 3.0 in each population (black) and those within each population that overlap with the UNL population (grey). Distribution of the iHS scores compared by **C**. Histograms and **D**. QQ plots.

**Additional file 5:** Table S2.xlsx. List of SNPs with Ensembl annotation of the nearest protein coding gene and iHS > -log p 3.0. It contains sheets for the TrypanoGEN populations and combined sheet for all population genes and DAVID derived Genetic associated diseases (GAD) and KEGG pathway affiliation

**Additional file 6:** Table S3.xlsx. List of signature SNPs associated with protein coding genes, with _iHS > -log p 3.0 and unique to the Nilo-Saharan Lugbara population.

**Additional file 7:** Figure S5.pdf. PANTHER Gene ontology database classification of the Nilo-Saharan Lugbara unique genes based on the Biological process and protein class they belong to. X-axis indicates the process and Y-axis the number of genes corresponding to the process.

**Additional file 8:** Table S4.xlsx. List of SNPs that are highly differentiated between the Niger-Congo and Nilo-Saharan Lugbara populations.

**Additional file 9:** Figure S6.pdf. Manhattan plot of the Rsb (**A**) and F_ST_ (**B**) against the whole genome (chromosome 1-22) between the Nilo-Saharan Lugbara (UNL) and TrypanoGEN Niger-Congo populations. **C**. Scatter plot of inter-population F_ST_ against Rsb scores for the SNPs having a high F_ST_ (> 0.99 percentile) and high Rsb (log p-value > 3.0) between UGN and the Niger-Congo TrypanoGEN populations. **D**. Venn diagram showing the number of intersecting protein coding SNP loci between the high Rbs-Fst (UNL verses All populations (CAM, CIV, GAS, UBB, ZAM) and high iHS (-log p-value > 3) protein coding loci unique to the UNL population only.

**Additional file 10:** Figure S7. Analysis of SNP loci in APOBEC3G gene (Chr22:39480259-39485103). **A**. Plot of SNP minor allele frequency (MAF) against chromosome position in the gene, the red dot is the position of rs112077004. **B**.Haploview plot of Linkage Disequilibrium (LD) of haplotypes within the gene, **C**. Zoomed LD plot of the SNPs upstream and downstream of the rs112077004 loci which is unique to the UGN population. **D**. Ensemble gene analysis of *APOBEC3G* gene locus on the GRCH37 build indicating the position of SNP rs112077004.

**Additional file 11:** Table S5.xlsx. Analysis of SNPs in the APOBEC3G gene including the minor allele frequency (MAF) and linkage disequilibrium (LD) statistics.

**Additional file 12:** Figure S8.pdf. Bar plots comparing the mean iHS *P*-values of the SNP loci in *APOL1*, *HPR*, *HLAG*, and *IL6* genes within the TrypanoGEN populations. These genes are associated with progression of Human African Trypanosomiasis.

**Additional file 13:** Table S6.xlsx. Population iHS scores of SNPs in genes that play a role in African trypanosomiasis pathology

